# A hierarchical point process model for spatial capture-recapture data

**DOI:** 10.1101/2020.10.06.325035

**Authors:** Wei Zhang, Joseph D. Chipperfield, Janine B. Illian, Pierre Dupont, Cyril Milleret, Perry de Valpine, Richard Bischof

**Affiliations:** Department of Environmental Science, Policy and Management, University of California Berkeley, Berkeley, USA; Faculty of Life Sciences and Natural Resource Management, Norwegian University of Life Sciences, Høgskoleveien 12, Ås, Norway; Norwegian Institute for Nature Research, Hyteknologisenteret, Thormøhlensgate 55, Bergen, Norway; School of Mathematics and Statistics, University of Glasgow, Glasgow, UK

**Author notes:** Correspondence Wei Zhang, Department of Environmental Science, Policy and Management, University of California, Berkeley, 130 Mulford Hall, Berkeley, CA, USA. The authors contributed equally to the manuscript (Wei Zhang, Joseph D. Chipperfield).

**Keywords:** Area search, Binomial point process, Continuous sampling, NIMBLE, Non-invasive genetic sampling, Poisson point process, Spatial capture-recapture, Wolverine

## Abstract

1. Spatial capture-recapture (SCR) is a popular method for estimating the abundance and density of wildlife populations. A standard SCR model consists of two sub-models: one for the activity centers of individuals and the other for the detections of each individual conditional on its activity center. So far, the detection sub-model of most SCR models is designed for sampling situations where fixed trap arrays are used to detect individuals.
2. Non-invasive genetic sampling (NGS) is widely applied in SCR studies. Using NGS methods, one often searches the study area for potential sources of DNA such as hairs and faeces, and records the locations of these samples. To analyse such data with SCR models, investigators usually impose an artificial detector grid and project detections to the nearest detector. However, there is a trade-off between the computational efficiency (fewer detectors) and the spatial accuracy (more detectors) when using this method.
3. Here, we propose a point process model for the detection process of SCR studies using NGS. The model better reflects the spatially continuous detection process and allows all spatial information in the data to be used without approximation error. As in many SCR models, we also use a point process model for the activity centers of individuals. The resulting hierarchical point process model enables estimation of total population size without imputing unobserved individuals via data augmentation, which can be computationally cumbersome. We write custom distributions for those spatial point processes and fit the SCR model in a Bayesian framework using Markov chain Monte Carlo in the R package nimble.
4. Simulations indicate good performance of the proposed model for parameter estimation. We demonstrate the application of the model in a real-life scenario by fitting it to NGS data of female wolverines (*Gulo gulo*) collected in three counties of Norway during the winter of 2018/19. Our model estimates that the density of female wolverines is 9.53 (95% CI: 8–11) per 10,000km^2^ in the study area.

## Introduction

Capture-recapture has a long history for estimating demographic parameters of wildlife populations such as abundance, survival rate, and density or relative abundance, which play a key role in wildlife conservation and management. Spatial capture-recapture (SCR), an extension to capture-recapture, has enjoyed rapid growth in the past decade. SCR models (see Borchers & Fewster, 2016, for a review) extend traditional capture-recapture models by incorporating the activity centers of individuals into the modelling framework as latent variables. The probability that an individual is detected by a detector (e.g., a trap) depends on the distance between the activity center of the individual and the detector. SCR models can therefore estimate spatially-explicit abundance of a population.

Standard SCR models are hierarchical and comprise two components: one for modelling the activity centers of individuals, and the other for modelling the detections of each individual conditional on its activity center. It is common to model the activity centers by a spatial point process since Efford (2004) used one for the first SCR model; see Chandler & Royle (2013); Dorazio (2013); Borchers *et al*. (2014) and Borchers & Marques (2017) for examples. A spatial point process describes the distribution of points in space, with both the number and locations of points being random. These models have been widely used for analysing spatial data in diverse fields such as histology, epidemiology, and seismology among many others (see Baddeley *et al*., 2006).

The detection model in SCR depends on the type of detectors used for data collection. Most detection models to date were developed for sampling situations in which the set of possible detection locations is fixed. This is the case when the data are collected using restraining devices such as live traps (see Powell & Proulx, 2003, for examples) or non-invasive techniques such as camera traps (Trolliet *et al*., 2014; Glover-Kapfer *et al*., 2019), hair snares (Henry *et al*., 2011), and acoustic arrays (Thomas & Marques, 2012). Other detection models have also been proposed for cases where detections are not necessarily associated with fixed locations. For example, Royle & Young (2008) proposed a SCR model to investigate temporary emigration of individuals. In their case, data are collected by an exhaustive area-search for individuals within a delineated sample unit, thus individuals can be detected anywhere in the unit. Similarly, Royle *et al*. (2011a) developed a model for ‘search-encounter’ SCR data that are collected by surveyors searching along transects.

Despite their variety (Royle *et al*., 2014; Efford, 2020), existing detector-based or search-encounter models do not adequately cover all common SCR sampling processes. Non-invasive genetic sampling (NGS) is now commonly used for collecting SCR data (Burgar *et al*., 2018; Ferreira *et al*., 2018; López-Bao *et al*., 2018). NGS in SCR studies involves the collection of DNA sources such as shed hair, faeces, saliva, and urine at fixed detector or area searches, followed by individual identification (Lamb *et al*., 2019). When NGS is implemented by searching a given area or along transects, paths taken by searchers can be recorded and used as a direct measure of effort in space and time. In praxis, this is not always possible, practical, or reliable. The model described in this manuscript is motivated by our experiences with the Scandinavian large carnivore monitoring program (Bischof *et al*., 2019), one of the world’s most extensive NGS projects (as of December 2019: over 35,000 samples from 6,000 individual bears (*Ursus arctos*), wolves (*Canis lupus*), and wolverines). Although field protocols prescribe the recording of GPS tracks during the performance of NGS by Norwegian and Swedish government employees, due to technical limitations and logistic constraints, these cannot always be linked unambiguously with detections. Furthermore, the region monitored is vast (over 500,000 km^2^) and a substantial portion of genetic samples are obtained via opportunistic detections by members of the public (hunters, hikers, berry pickers, etc.). As a consequence, no direct measure of effort is available and investigators wanting to account for spatial heterogeneity in detection probability are left with proxies for effort such as distance to roads and snow cover (Bischof *et al*., 2019). In cases such as this, detections, at least in theory, are possible at any location within the general area that humans could visit. An approach that uses actual detection locations would be preferable to the typical approach of projecting detections to an artificial detection grid (Russell *et al*., 2012; Lonsinger *et al*., 2018; López-Bao *et al*., 2018) as the latter: (a) means an approximation of detection locations, (b) involves aggregation of detection information, and (c) forces investigators to trade off precision for computational efficiency (Milleret *et al*., 2018).

To deal with SCR data collected through area searches, we propose a hierarchical point process modelling framework. As in most SCR models in the literature, we model the activity centers of individuals by a spatial point process, which in our case is an inhomogenous Poisson point process. We assume these activity centers do not change during the study, although it is possible to extend the framework proposed here to include movement. A second Poisson point process is used to model the detections of an individual conditional on its activity center.

A Poisson point process is determined by an intensity function. Evaluating the intensity function at point ***x*** gives the probability that there is one point (one detection or activity center in our case) falling within an infinitesimal volume centered around ***x*** (p.51, Chiu *et al*., 2013). The detection intensity for one individual at one point in the study area is a (typically decreasing) function of the distance between the individual’s activity center and that point. The expected number of detections for one individual in the study area or any subregion is obtained by integrating the intensity function over the area or subregion. By eliminating the need for an artificial grid of detectors, the point process model represents more closely the spatially-continuous data collection process. All spatial information in the data can be used without additional post-collection approximation error. We fit the proposed hierarchical model in a Bayesian framework using Markov chain Monte Carlo (MCMC).

Estimating total population size when fitting SCR models using MCMC leads to technical challenges because it involves estimating the number of individuals that were completely unobserved. Most authors do this using data augmentation (Royle *et al*., 2007; Royle & Dorazio, 2012), in which unobserved individuals, their activity centers, and their state (real or virtual) are imputed as part of the MCMC posterior sampling. This can be computationally costly, especially when detection rates are low and hence there may be many unobserved individuals. A more efficient solution is to construct a ‘semi-complete data likelihood’, which is the product of a complete data likelihood of observed individuals and a marginal likelihood of unoberved individuals obtained by integrating out their activity centers (King *et al*., 2016). Then, posterior sampling for model parameters and the activity centers of observed individuals can be achieved using MCMC and the semi-complete data likelihood without data augmentation. This is achieved by deriving the marginal detection probability, which is calculated by numerical integration.

We investigate performance of the proposed model and estimation method for recovering parameters using a simulation study. We also apply the model to analysis of NGS data of female wolverines collected during the winter of 2018/19 in three counties of Norway. Due to the spatial scale of the study area and the nature of the detection sampling, these data are difficult to analyze with other approaches for modelling detection.

## Materials and methods

### Notation

A set of notation used in following sections is summarized here:

- *N* : the population size.
- *N* *: the number of individuals detected at least once.
- *D*: the dimension of the spatial domain (study area): *D* = 2.
- *M*_*i*_: the observed number of detections for individual *i* = 1, …, *N*.
- *D*_*i*,***o***_: the number of detections of individual *i* across the entire region ***o***.
- ***õ***: the entire region for the placement of activity centers (habitat selection).
- ***o***: the entire region where individuals can be detected; ***o*** may or may not be the same as ***õ***.
- 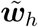: a sub-window of ***õ***; *h* = 1, …, *H*.
- ***w***_*l*_: a sub-window of ***o***; *l* = 1, …, *L*.
- ***s***_*i*_: location of the activity center of individual *i*; ***s***_*i*_ *∈* ***õ***; *i* = 1, …, *N*.
- ***d***_*i,j*_: location of the *j*-th detection for individual *i*; ***d***_*i,j*_ *∈* ***o***; *i* = 1, …, *N, j* = 1, …, *M*_*i*_.
- ***β***: a vector of parameters that determine the intensity function of the point process model for habitat selection.
- ***θ***: a vector of parameters that determine the baseline detection function.
- ***σ***: a vector of parameters that determine the (general) detection kernel function.
- *σ*: a scalar parameter that determines the (specific) isotropic multivariate Gaussian detection kernel function, which is given in Appendix II.
- 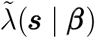: the intensity function of the point process model for habitat selection.
- *λ*(***d*** | *s*, ***θ, σ***): the intensity function of the point process model for detection.
- 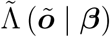:the expected number of activity centers within the region ***õ***.
- Λ (***o*** | ***s***_*i*_, ***θ, σ***): the expected number of detections of individual *i* within the region ***o***.
- 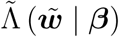: the expected number of activity centers within sub-window 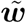.
- Λ (***w*** | ***s***_*i*_, ***θ, σ***): the expected number of detections of individual *i* within sub-window ***w***.

For convenience, we use ⎰_**Ω**_ *f* (***x***)*d****x*** to denote the multi-dimensional integral of *f* (***x***) over the region **Ω**, which can be ***o, ***õ***, w*** or 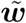.

### Habitat selection

The activity center of an individual represents the center of the individual’s home range, around which the individual moves. Spatial point processes have been widely used for modelling the placement of activity centers since Efford (2004); see Illian *et al*. (2007) for an introduction to spatial point processes. In most studies, such as Chandler & Royle (2013), a homogenous Poisson point process is used, whose intensity function 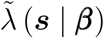is a constant across ***õ***, the entire region of interest. Although less widely used, inhomogenous Poisson point processes have also gained some applications in modelling habitat selection (e.g., Dorazio, 2013; Borchers *et al*., 2014; Borchers & Marques, 2017). Using an inhomogeneous Poisson process, it is possible to model a spatially varying intensity surface, which allows for a higher density of activity centers in areas with higher habitat quality, and to study second-order habitat selection (Johnson, 1980).

Following Illian *et al*. (2007, p.121), the log-probability density of there being *N* activity centers, ***s***_1_, …, ***s***_*N*_, over the entire region ***õ*** is

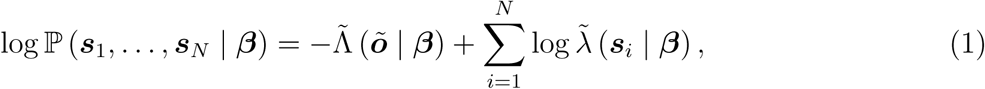

Where

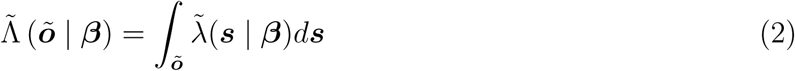

is the expected number of activity centers over ***õ***.

### Detection intensity

Key to formulating a model for the detection process in SCR is to construct a detection intensity function, which may be related to the observed data in different ways. For example, if one individual can be detected by one detector multiple times, then the number of detections for the individual at that detector can be modelled by a Poisson distribution with mean equal to the intensity (see López-Bao *et al*., 2018, for example). In cases where an individual can only be detected once by a given detector during one sampling period, the intensity can be defined as the probability of detecting the individual at that detector. The detection outcome for the individual at that detector follows a Bernoulli distribution (see Royle *et al*., 2011b, for example). In addition, to improve computational efficiency the Bernoulli detection model can be adequately approximated by a binomial model with the detection locations aggregated at coarser spatial scales (Milleret *et al*., 2018).

For most current SCR models and the model we develop below, the detection intensity is of the following form

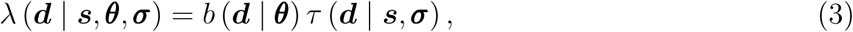

where ***d*** denotes the location of the detection, ***s*** denotes the activity center, *τ* (***d*** | ***s, σ***) is a kernel describing the distance decay relationship of the detection intensity from the activity center, and *b* (***d*** | ***θ***) is a function describing the baseline detection intensity at the detection location. The function *τ* (***d*** | ***s, σ***) describes the decreasing chance of detecting an individual as the distance from the activity center increases and is therefore a functional description of the home range of the individual. The function *b* (***d*** | ***θ***) represents the heterogeneity in detection probability that might result from differences in one or more aspects of the following: (1) sampling effort, (2) the effects of the landscape on individual detectability, and (3) trap efficiency such as the type of camera traps (Efford *et al*., 2013).

### Spatial point process model for detection

When SCR data are collected by searching the study area, the location of each detection can be any point in ***o***. Here, we model the detections of each individual using a spatial point process. A class of spatial point process models that serves as an initial framework for this purpose is the inhomogenous Poisson process.

The log-probability density for the *M*_*i*_ detection locations, 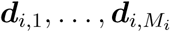, of *individual i given that* its activity center is ***s***_*i*_ is:

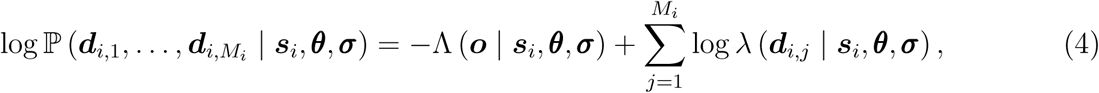

Where

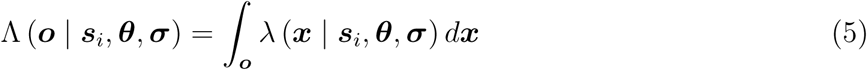

is the expected number of detections of individual *i* over the entire detection region. We assume that detections of each of the *N* individuals are conditionally independent given their respective activity centers. It follows that the log-probability density for the detections of all individuals in the population is the sum of the individual log-probability densities:

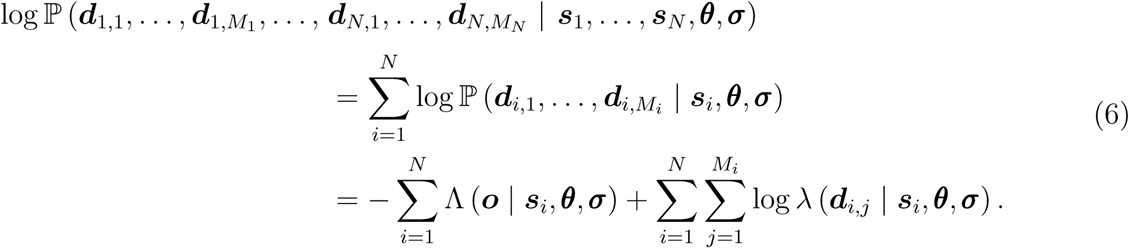

### Marginal void probability

In all SCR studies, there are some individuals in the population that are never detected. The probability of an individual in the population never being detected throughout the study plays a key role in formulating our hierarchical modelling framework below. For spatial point processes, this is related to the so-called ‘void probability’, the probability that no points fall within a given region. From Equation (25), the void probability ℙ (*D*_*i,o*_ = 0 | ***s***_*i*_, ***θ, σ***) for individual *i* is

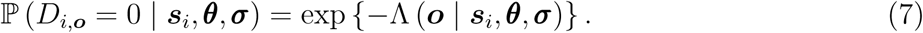

The void probability depends on the unknown activity center ***s***_*i*_ of individual *i*, so it cannot be used for our purpose directly. We need the marginal void probability, which is derived below.

A known result of the Poisson point process is that conditioning on the total number of points in a given region yields a binomial point process (p.69, Illian *et al*., 2007), a point process with a fixed number of points. For a binomial point process, the probability density of one point’s location is the intensity of the corresponding Poisson process evaluated at that point divided by the integral of the intensity function over the entire region. It follows that given individual *i* exists in the population, the probability density that it has activity center ***s***_*i*_ is

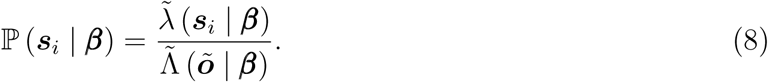

Thus, given there are *N* activity centers in the region of interest ***õ***, the log-probability density for activity centers ***s***_1_, …, ***s***_*N*_ is

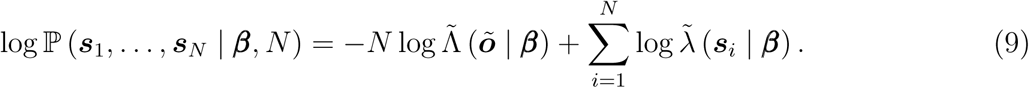

From Equations (7) and (8), it follows that the marginal void probability *p*\* that individual *i* is never detected is

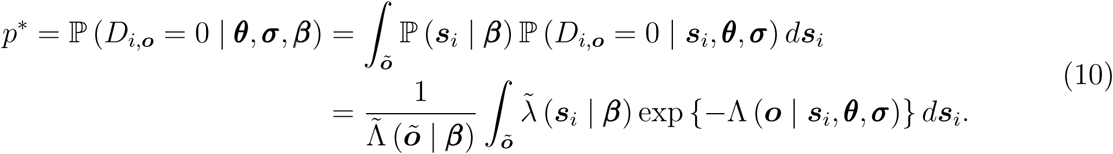

It is immediate that the marginal void probability is the same for all individuals in the population. Calculating *p*\* seems complicated; however, for many commonly-used habitat selection and detection intensity functions, it is possible to express the formula for calculating *p*\* either in terms of elementary functions or non-elementary functions with computationally tractable numerical approximations. One example of the Gaussian kernel function is given in the Appendix.

### Hierarchical point process model

Combining the models for habitat selection and detection described above results in a hierarchical point process SCR model. As shown in Figure [No.], the full model consists of four parts:

- The activity centers of individuals are modelled by a Poisson point process with intensity 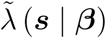. Then the population size *N* follows a Poisson distribution:

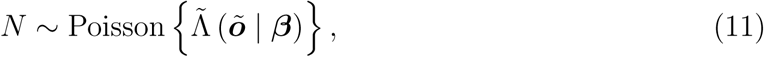

where the expected population size 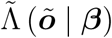 is given in Equation (2).
- Conditional on population size *N*, the number of individuals detected, *N* *, follows a binomial distribution:

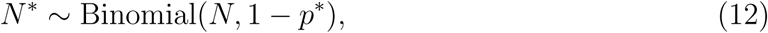

where *p*\* is the marginal void probability given in (10).
- Conditional on the number of detected individuals *N* *, the activity centers of the detected individuals, ***s***_1_, …, ***s***_*N\**_, follow a binomial point process, whose intensity function is given in (8).
- Conditional on the activity center ***s***_*i*_ of individual *i* = 1, …, *N* *, the detections 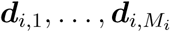 follow a Poisson point process with intensity *λ* (***d*** | ***s***_*i*_, ***θ, σ***). The log-probability density for these detections is:

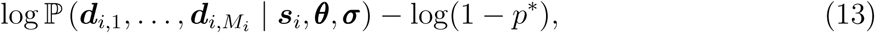

where 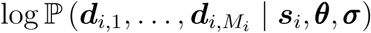 is given in Equation (4). Note that Equation (13) contains the term − log(1−*p*\*) since here we condition on the fact that the individual is detected, which has probability 1 − *p*\*.

### Cell-based framework for covariates

SCR models often incorporate covariates to improve the accuracy of inferences. For example, available resources are usually considered when evaluating how animals distribute themselves in space (Gaillard *et al*., 2010). Using our point process model, it is straightforward to use covariates for either the habitat selection process or the detection process. This can be achieved by formulating the intensity of the inhomogeneous Poisson process as a function of covariates.

While in some cases it is possible to derive the environmental covariates as spatial functions, it is more common for environmental data to be retrieved, processed, and distributed as cell-based discrete approximations to the continuous process. To handle such covariates for habitat selection, the entire region of interest ***õ*** is divided into a set of *H* non-overlapping habitat sub-windows 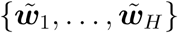 such that 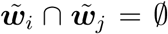 if *i* ≠*j* and 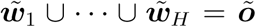. Similarly, for cell-based detection covariates we divide the detection region **o** into a set of of *L* non-overlapping detection sub-windows ***w***_1_, …, ***w***_*L*_ such that ***w***_*i*_ *∩* ***w***_*j*_ = *∅* if *i* ≠ *j* and ***w***_1_ *∪ … ∪* ***w***_*L*_ = ***o***. The value of each covariate is constant within each sub-window and is related to the intensity value of the sub-window through some link function. In principal, any non-negative function is valid for this purpose. Here, we consider the log-linear model as an example for both habitat selection and detection. Define 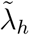 to be the habitat selection intensity value of sub-window 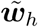, *h* = 1, …, *H*. It follows that

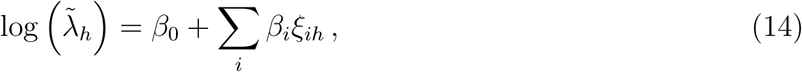

where *ξ*_*ih*_ is the value of the *i*-th habitat selection covariate in sub-window 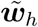. Similarly, define *b*_*l*_ to be the baseline detection intensity of sub-window ***w***_*l*_, *l* = 1, …, *L*. Then we have

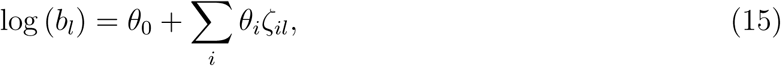

where *ζ*_*il*_ is the value of the *i*-th detection covariate within sub-window ***w***_*l*_.

In situations where orthogonally neighbouring sub-windows share the same values for the covariates, it may be possible to merge these sub-windows into one. This minimises the number of sub-windows and thus reduces the amount of computation needed for model fitting. The size of the habitat and detection sub-windows should be chosen according to computational considerations but also at the scale relevant to the ecological and observation processes (Johnson, 1980; Gaillard *et al*., 2010).

Using the grid cells above, we can rewrite the intensity functions of the Poisson point processes for habitat selection and detection as two piecewise constant functions; see Appendix I and II for details. This reduces the calculation of the definite integrals in (2) and (5) to discrete summations. As a result, calculating the marginal void probability (10) is simplified to some extend; see Appendix III.

### Software implementation

We fit the model via Bayesian MCMC in the R package nimble (de Valpine *et al*., 2017). nimble supports nearly the same modelling language as JAGS and WinBUGS but allows extensions with new functions and distributions. We used these capabilities to write both Poisson and binomial point process distributions and also marginal void probability calculations in nimble, allowing them to be automatically incorporated into MCMC sampling.

## Results

### Simulation study

We designed a simulation study to investigate the relative bias and frequentist coverage of credible intervals for calibrated Bayes interpretation (Little, 2006) under a range of covariate effects. We set the habitat region ***õ*** to be a 10km×10km square, which included a buffer of 0.6km around the detection region ***o***, a 8.8km×8.8km square. The region ***õ*** was divided into 100 equally-sized sub-windows and the region ***o*** was divided into 25 equally-sized sub-windows. We considered two spatial covariates in the simulations, one for modelling the habitat selection intensity and the other for the baseline detection intensity. Values of both covariates were simulated from a uniform distribution: Uniform(−1, 1). The intercept parameters *β*_0_ and *θ*_0_ were set to be 1.0 and 2.0 respectively. The values of the slope parameters *β*_1_ and *θ*_1_ were chosen from set *{*−1, 0, 1*}* so that there were nine scenarios defined by different combinations for the two parameters. An isotropic multivariate Gaussian function with *σ* = 0.2 was used for the detection kernel.

For each of the nine scenarios, we simulated 100 datasets under the hierarchical point process model. The model was then fit to the data using Bayesian MCMC in nimble. MCMC sampling was run for 110,000 iterations with the first 10,000 discarded as burn-in. Estimation results of the population size *N* are shown in Fig. 1. It can be seen that almost unbiased inference with roughly a nominal value for the credible interval coverage was obtained for all the simulations. Similar conclusions can be drawn from the results of other parameters shown in the supplementary material. We therefore conclude that the proposed model works well for parameter estimation.

**Figure 1:**
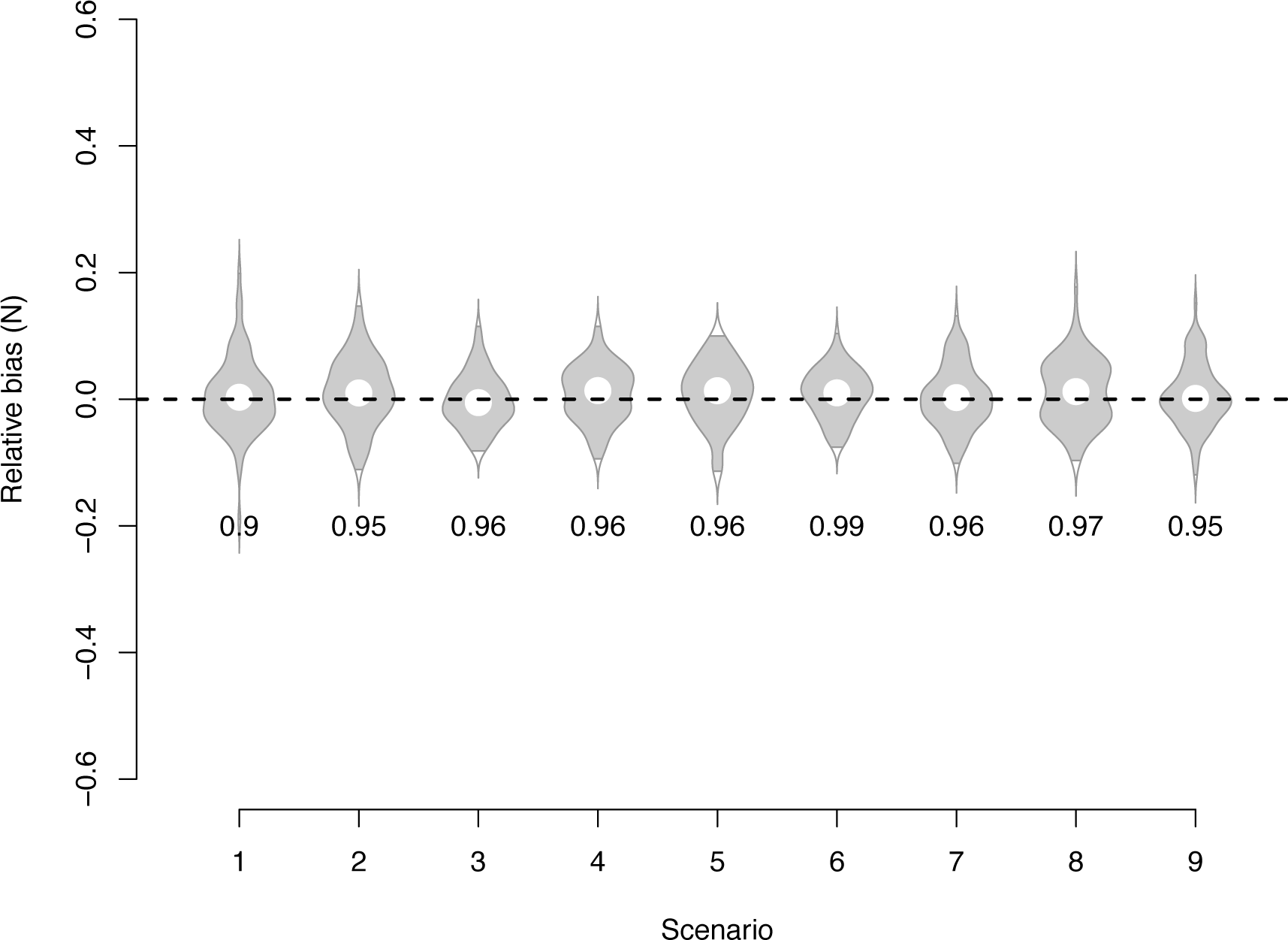
Violin plots showing the distribution of relative estimation error (defined as (estimate − true value) / true value) of the population size *N* based on 100 replicates for each scenario. The value of (*β*_1_, *θ*_1_) in scenarios 1–9 is (− 1, − 1), (0, − 1), (1, − 1), (− 1, 0), (0, 0), (1, 0), (− 1, 1), (0, 1), and (1, 1) respectively. The value below each violin plot gives the credible interval coverage.

### Wolverine data analysis

To demonstrate the application of the proposed model, we applied it to part of the wolverines data available in the Scandinavian large carnivore database, Rovbase 3.0 (http://rovbase.se/orhttp://rovbase.no/). This database is jointly used by Norway and Sweden to record NGS data, dead recoveries, GPS search tracks, and observations of wolverines and other large carnivores. Here, we restricted our analysis to the data of female wolverines collected from December 2018 to June 2019 in Norwegian counties of Hedmark, Oppland, and parts of Sør-Trøndelag (Fig. 2; Flagstad et al. 2004, Brøseth et al. 2010, Gervasi et al. 2015). The dataset is composed of 228 scat- and hair-based DNA samples from 72 female wolverines.

**Figure 2:**
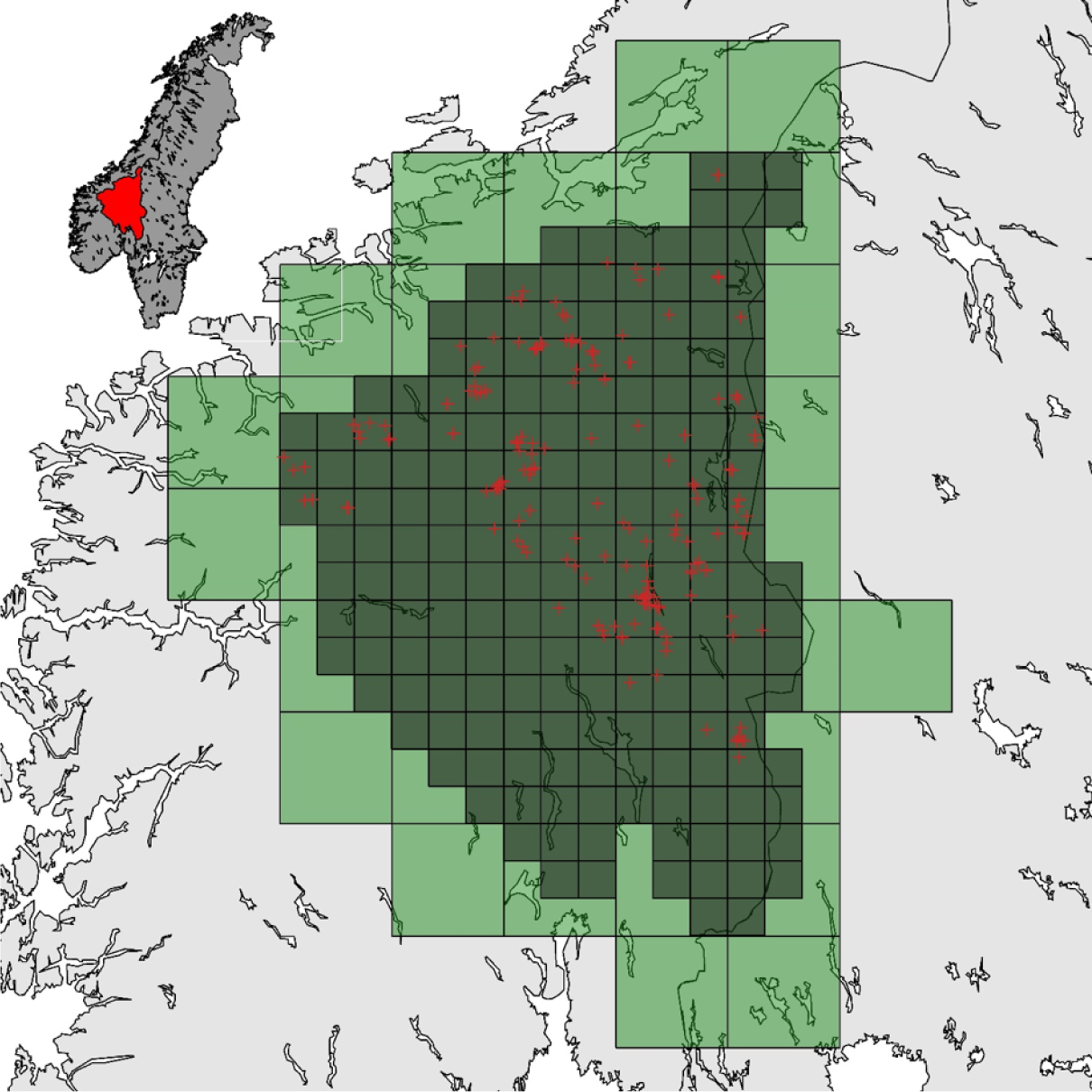
Distribution of non-invasive DNA samples (red crosses) of female wolverines collected during winter 2018/19 in Norway. The area that was searched for samples is represented by the red polygon on the map in the upper left corner. Detection sub-windows (20km×20km each) are represented by the dark green grid cells. Habitat selection sub-windows (60km×60km each) are represented by the larger light green grid cells.

As shown in Fig. 2, the detection region ***o*** was divided into *L* = 195 sub-windows each of size 20km×20km and the habitat selection region ***õ*** was divided into *H* = 40 sub-windows each of size 60km×60km. The region ***õ*** covers the entire region ***o*** and a surrounding buffer, allowing the activity centers of individuals to be located outside the searched area. The number of known wolverine dens was used as a covariate for modelling habitat selection. Five covariates *ζ*_*i*_, *i* = 1, …, 5 were used for modelling baseline detection. Indicator variables *ζ*_1_ and *ζ*_2_ take value 1 if the detection location is in Oppland and Sør-Trøndelag, respectively, and take value 0 otherwise. In addtion, covariates *ζ*_3_, *ζ*_4_, and *ζ*_5_ denote respectively the recorded length of GPS tracks logged by searchers, the average percentage of snow cover (MODIS at 0.1 degrees resolution, www.neo.sci.gsfc.nasa.gov, accessed 2019-10-11), and the average distance to the nearest primary and secondary roads. The isotropic multivariate Gaussian kernel function was used for detection

To fit the data, we ran eight MCMC chains for 20,000 iterations and removed the first 5,000 iterations as burn-in. The model was assumed to have reached convergence when all R-hat values were *≤* 1.1 and visual inspections of trace plots revealed good mixing (Brooks & Gelman, 1998). Mean parameter estimates, standard deviations, 95% credible intervals, R-hat values, and effective sample sizes are shown in Table 1. It is estimated that on average there are 9.53 (95% CI: 8–11) female wolverines per 10,000 km^2^ within the sampled area.

**Table 1:** Estimation results for the hierarchical point process model fitted to the female wolverine data collected in eastern Norway during winter 2018/2019. ESS represents the effective sample size.

| Parameters | Mean | SD | 2.5% | 97.5% | R-hat | ESS |
| --- | --- | --- | --- | --- | --- | --- |
| $N$ | 137.19 | 12.25 | 115 | 163 | $< 1.001$ | 5,761 |
| $\beta_0$ | -21.04 | 0.15 | -21.35 | -20.76 | 1.002 | 2,626 |
| $\beta_1$ | 0.64 | 0.11 | 0.42 | 0.86 | $< 1.001$ | 5,008 |
| $\theta_0$ | -18.38 | 0.83 | -20.05 | -16.79 | $< 1.001$ | 13,771 |
| $\theta_1$ | -18.28 | 0.21 | -18.71 | -17.88 | $< 1.001$ | 12,538 |
| $\theta_2$ | -17.99 | 0.14 | -18.27 | -17.71 | 1.002 | 2,980 |
| $\theta_3$ | 0.25 | 0.09 | 0.08 | 0.43 | $< 1.001$ | 5,120 |
| $\theta_4$ | -0.13 | 0.09 | -0.31 | 0.05 | $< 1.001$ | 11,540 |
| $\theta_5$ | 0.14 | 0.10 | -0.06 | 0.34 | $< 1.001$ | 12,452 |
| $\sigma$ | 5.16 | 0.21 | 4.76 | 5.59 | 1.001 | 5,121 |

## Discussion

We developed a hierarchical point process model for SCR data collected by exhaustive area searches. The development of the model was motivated by the NGS data of large carnivores in Scandinavia, where individuals can, at least in theory, be detected anywhere within the sampled area. Fitting such data via the artificial detector grid approach leads to spatial information loss and computational efficiency issues (Milleret *et al*., 2018). The search-encounter model of Royle *et al*. (2011a) is not suitable in this context because auxiliary information (i.e. sampling paths) on sampling effort is not completely available.

Our model is a reasonable reflection of the NGS data collection process and all spatial information in the data can be used without additional approximation error. Spatial covariates can be incorporated to model the intensity functions of the inhomogeneous Poisson point processes for both habitat selection and detection. In addition, we formulated the hierarchical model using the void probability approach, which makes it possible to fit the model via MCMC in a Bayesian paradigm without needing to use data augmentation. This improves the efficiency of model fitting and eliminates the implementation challenges associated with data augmentation. We demonstrated the promising performance of the proposed model for parameter estimation with a set of simulations and the wolverine data analysis.

The approach presented here can be extended to work for several more advanced SCR models. First, we could extend the modelling framework to open populations by adding an extra layer to the current model to account for the changes in population size and structure over time. It follows that the population size *N* will be conditional on the population model used rather than the intensity function for habitat selection. The distribution of activity centers then becomes a binomial point process instead of Poisson. In this case, 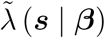 becomes a function describing the relative density of activity centers rather than the absolute density. Second, SCR data with group identities (e.g., Chandler & Royle, 2013; Augustine *et al*., 2018) can be handled easily using our point process detection model. Suppose that there are two sets of spatial points, each of which is generated from a Poisson point process. Then points in the union of the two sets follow another Poisson point process, whose intensity is the sum of the intensities of the two separate Poisson processes (Cinlar & Agnew, 1968). This means that detections from a group of non-uniquely-identifiable individuals can be modelled as a single Poisson point process. It is immediate that the framework presented in this manuscript can be applied directly to SCR data with group identities.

## Acknowledgements

The study was funded by the Norwegian Environment Agency (Miljødirektoratet), the Swedish Environmental Protection Agency (Naturvårdsverket), the Research Council of Norway (NFR 286886; project WildMap), and the Peder Sather Grant. We thank Henrik Brøseth for comments on the manuscript.

## Authors’ contributions

JDC, PD, CM, and RB conceived the ideas with input from JBI and PdV. WZ, JDC, and JBI formulated the mathematical framework of the model. WZ, JDC, and PdV implemented the model in NIMBLE. PD, CM, and RB performed the analysis of the wolverines data. WZ and JDC led the writing of the manuscript. All authors contributed critically to the drafts and gave final approval for publication.

## Data availability

The NGS wolverine data and R code used in this manuscript will be available on GitHub (https://github.com/wzhang721) after the manuscript is accepted.

## APPENDIX I: cell-based habitat selection

Note that 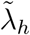 is the habitat selection intensity value of sub-window 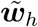, *h* = 1, …, *H*. It follows that the habitat selection intensity function 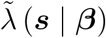 can be written as

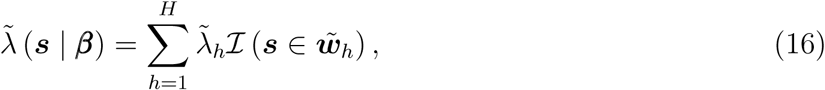

where *ℐ* (·) is the usual indicator function that equals 1 if the condition · is true and 0 otherwise.

Provided that an inhomogenous Poisson point process is used for habitat selection, the number of activity centers 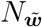 within a subregion 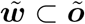 follows a Poisson distribution:

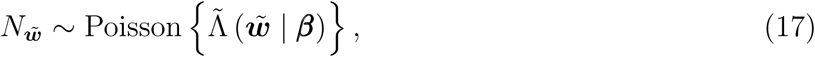

Where

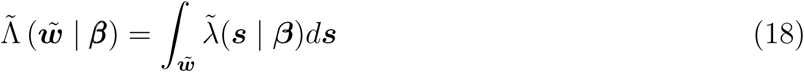

is the expected number of activity centers located within 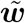. According to Equation (18), the expected number of activity centers within sub-window 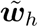 is

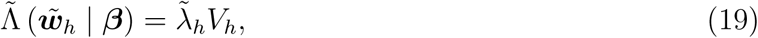

where *V*_*h*_ is the volume of sub-window 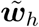.It follows that the expected total number of activity centers across the entire region ***õ*** is

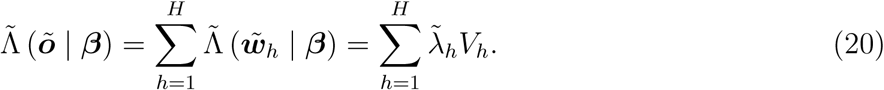

## Appendix II: cell-based detection

Note that *b*_*l*_ denotes the baseline detection intensity of sub-window ***w***_*l*_, *l* = 1, …, *L*. Then, similarly to the habitat selection case above, the baseline detection intensity function can be written as

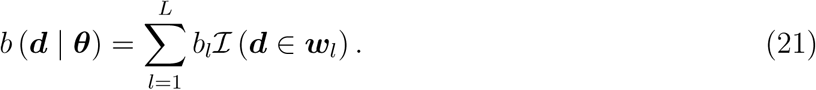

A common detection decay kernel is the isotropic multivariate Gaussian kernel

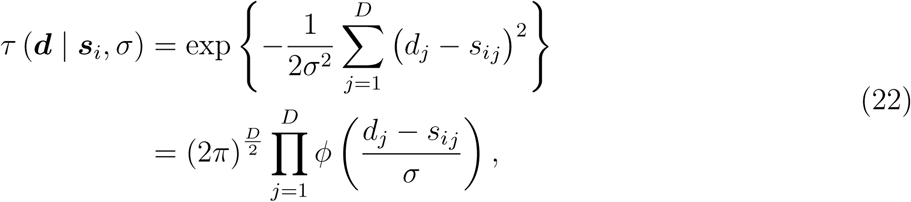

where *Φ* (*x*) is the standard normal probability density function

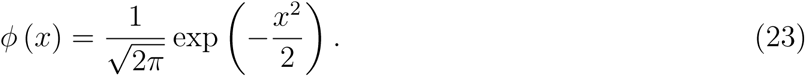

Here, *τ* (***d*** | ***s***_*i*_, *σ*) is proportional to the probability density function of a multivariate Gaussian distribution with mean equal to the activity centre ***s***_*i*_ of individual *i*, and with a diagonal variance-covariance matrix whose diagonal elements are all set to *σ*^2^, representing the isotropic variance of the detection decay kernel. Thus, the parameter *σ* regulates the home range size of the species of interest. In this paper, we do not consider covariates for *σ*. However, it is immediate that *σ* can be modelled as a regression of relevant environmental and/or individual covariates allowing for different home range sizes in different environments or for species that have different home ranges at different life stages or between different sexes.

Inserting the cell-based baseline detection function (21) and the Gaussian decay kernel (22) into Equation (3) yields the following composite detection intensity function

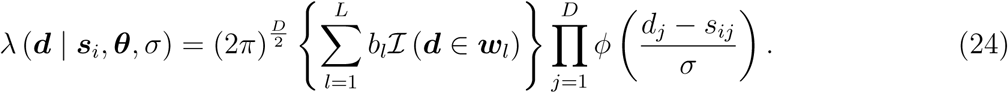

### Resulting marginal void probability

Similar to the point process for activity centers, the number of detections *D*_*i,w*_ of individual *I* within a subregion ***w*** *⊂* ***o*** follows a Poisson distribution:

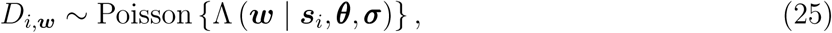

where

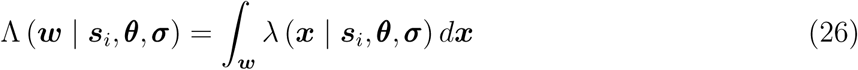

is the expected number of detections within ***w***. Then the expected number of detections Λ (***w***_*l*_ | ***s***_*i*_, ***θ***, *σ*) of individual *i* within sub-window ***w***_*l*_ is

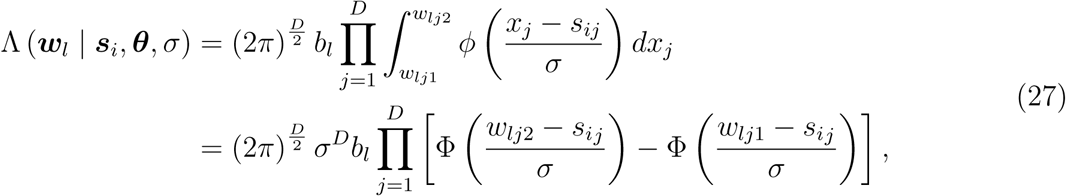

where *w*_*lj*1_ and *w*_*lj*2_ are the lower and upper bounds of the *j*-th dimension of sub-window ***w***_*l*_, and Φ (*x*) is the standard Gaussian cumulative distribution function. Then the expected number of detections of individual *i* across the entire region ***o*** is

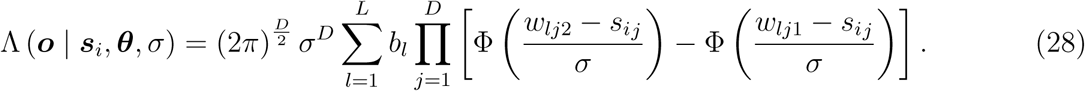

According to Equation (10), the marginal void probability in the case of cell-based habitat selection and detection is

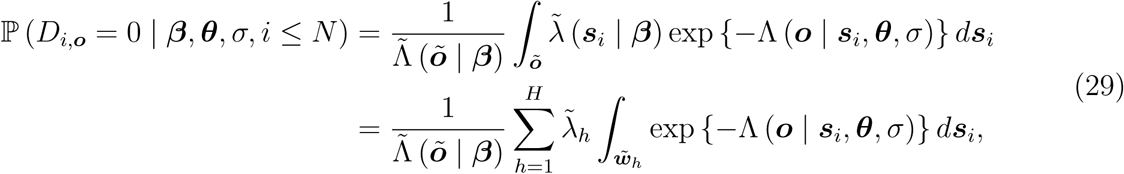

where 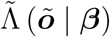 is given in Equation (20), and the definite integral 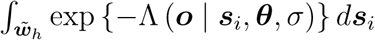 can be calculated using the grid integration of Equation (28).

## Notes

### Competing Interest Statement

The authors have declared no competing interest.

